# GeneF: A High-Performance Processing-in-Memory Accelerator for Efficient DNA Alignment

**DOI:** 10.1101/2025.08.08.669185

**Authors:** Kanchan Verandani

## Abstract

In this paper, we explore the compute and memory characteristics of the FM-index and identify data movement as a significant contributor to overall energy consumption in genomic processing. We propose GeneF, a Processing-in-Memory (PIM) accelerator designed specifically for DNA alignment tasks, leveraging 3D-stacked memory to enhance memory bandwidth and computing parallelism. Our architecture features a custom RISC-V-based processing element (PE) array, a lightweight messaging mechanism to mitigate remote access latency, and specialized prefetchers for improved efficiency. Experimental results demonstrate that GeneF achieves substantial speedups—1820× for counting and 1728× for determining stages—over traditional CPU implementations and offers remarkable energy efficiency, consuming only 25% of the energy compared to conventional CPU-DDR3 systems. The findings highlight the potential of PIM architectures in minimizing data movement and enhancing performance for genomic workloads, paving the way for more energy-efficient computing solutions in the field of bioinformatics.

## I. Introduction

Precision medicine has emerged as a transformative approach in clinical diagnostics, enabling treatments tailored to the unique genetic makeup of individual patients. This paradigm shift towards personalized healthcare is especially impactful in oncology, where genetic profiling of tumors from cancers such as melanoma, leukemia, breast, lung, colon, and rectal cancers guides targeted therapies [1]–[3]. The increasing adoption of genomic sequencing in clinical settings is expected to generate an unprecedented volume of data. It is estimated that by 2025, over one billion individuals worldwide will have their genome sequenced, resulting in the production of approximately 40 exabytes of genomic data annually [4]. This explosive growth in genomic data demands the development of advanced computational tools capable of efficient and scalable analysis.

A fundamental step in processing genomic data is *sequence alignment*, where short DNA fragments (reads) generated by sequencing machines are mapped back to a reference genome. Most state-of-theart alignment algorithms follow a *seed-and-extend* paradigm. Initially, the *seed* step involves rapidly locating exact matches or candidate regions within the reference genome, often through an index structure. Subsequently, the *extend* step performs a more computationally intensive inexact matching, typically using dynamic programming methods such as the Smith-Waterman algorithm, to refine alignments based on scoring schemes.

The computational cost of these two stages varies significantly depending on the sequencing technology. Next-generation sequencing (NGS) technologies produce relatively short reads, typically ranging from 50 to 200 base pairs, which contrasts with the much longer reads generated by earlier or third-generation platforms. While SmithWaterman’s quadratic time complexity, 𝒪(*N* ^2^), where *N* is the read length, can be mitigated through hardware accelerators like Darwin coprocessors, these accelerators are optimized for long-read sequences (greater than 1,000 base pairs). Consequently, they do not efficiently accelerate NGS short-read alignment, as the *extend* step does not dominate the runtime for such reads.

In the case of widely used NGS alignment tools like BWA [5], the majority of computational time (ranging from 29% to 70%) is spent on the *FM-index search* within the *seed* phase rather than the *extend* phase. The FM-index is a compressed full-text substring index based on the Burrows-Wheeler transform that enables fast exact matching. Despite its importance, accelerating FM-index search remains challenging due to two primary factors: the irregular memory access patterns and the relatively lightweight computational operations involved.

This work explores the application of processing-in-memory (PIM) architectures as a promising solution to these challenges. PIM integrates computation capabilities directly within memory devices, effectively mitigating the bandwidth bottleneck caused by frequent data movement between main memory and processing units. Through a detailed analysis of the FM-index algorithm, we identify that the underlying operations are predominantly simple, such as bitwise and arithmetic computations, yet they generate substantial data traffic. These characteristics make FM-index highly amenable to PIM acceleration, where lightweight compute units embedded within memory can execute these operations in parallel, significantly reducing data movement overhead.

We propose **GeneF**, a novel, application-driven accelerator designed to exploit PIM for FM-index search. Our design addresses the stringent constraints of processing elements (PEs) in PIM environments by maintaining simple but efficient computation units. GeneF leverages the massive inherent parallelism in FM-index operations and introduces an efficient communication mechanism between PEs residing on the logic layer of PIM hardware. Additionally, we develop a specialized prefetcher that decouples data access from computation, enabling full pipelining and overlapping of memory accesses with computation to maximize throughput.

Our comprehensive evaluations demonstrate that GeneF significantly outperforms existing solutions. It achieves a 20-fold speedup over the best available ASIC accelerators tailored for FM-index, and compared to the widely used BWA-MEM software running on conventional hardware, GeneF delivers an impressive 1820× improvement in performance along with a 7032× enhancement in energy efficiency.

The key contributions of this paper can be summarized as follows:

- We identify data movement between memory and compute units as the dominant factor limiting FM-index performance. The algorithm’s reliance on simple, parallelizable operations motivates the use of lightweight PIM-based compute units.
- We introduce GeneF, an application-specific PIM accelerator for FM-index that features a scalable communication infrastructure among PEs and a dedicated prefetcher, facilitating efficient overlap of computation and data access.
- We provide an extensive evaluation showing that GeneF substantially outperforms current ASIC solutions and conventional software implementations, demonstrating the effectiveness of PIM for genomic data processing acceleration.

## II. Background and Motivation

### A. FM-index Algorithm Overview

The FM-index [6] is a foundational data structure widely adopted in numerous genomic alignment tools [7] due to its efficiency in exact pattern matching within large reference sequences. This indexing method aligns well with the *seed-and-extend* paradigm commonly used in sequence alignment workflows. During the *seeding* phase, short subsequences (seeds) extracted from the query reads are precisely matched back to the reference genome without allowing gaps or mismatches.

The construction of the FM-index relies on the Burrows-Wheeler Transform (BWT) of the reference sequence. For illustration, consider a simple genomic sequence *T* = “abracadabra$”. The BWT process involves the following steps:

1. Generate all cyclic rotations of the string *T*.
2. Sort these rotations lexicographically to form a matrix *M*.
3. Extract the last column *L* of this matrix, which constitutes the BWT string of *T*. For the example, the BWT of “abracadabra$” results in the string “ard$rcaaaabb”.

Two key properties underpin the FM-index’s ability to facilitate efficient search. First, the sorted rotations enable quick identification of substrings based on their lexicographical order. Second, the *last-to-first* mapping property ensures that the *i*^*th*^ occurrence of a character in the last column corresponds to the *i*^*th*^ occurrence of the same character in the first column. Formally, the last-to-first (LF) mapping function is defined as:

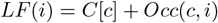

where *C*[*c*] denotes the count of all characters lexicographically smaller than *c* in the BWT, and *Occ*(*c, i*) is the number of occurrences of character *c* up to position *i* in the BWT string. This property enables backward search over the FM-index to efficiently locate all occurrences of a query substring in the reference.

The pseudocode for the FM-index based seeding algorithm is shown in Algorithm 1, which iteratively refines the search interval [*sp, ep*] by extending the query string from right to left.

**Algorithm 1:**
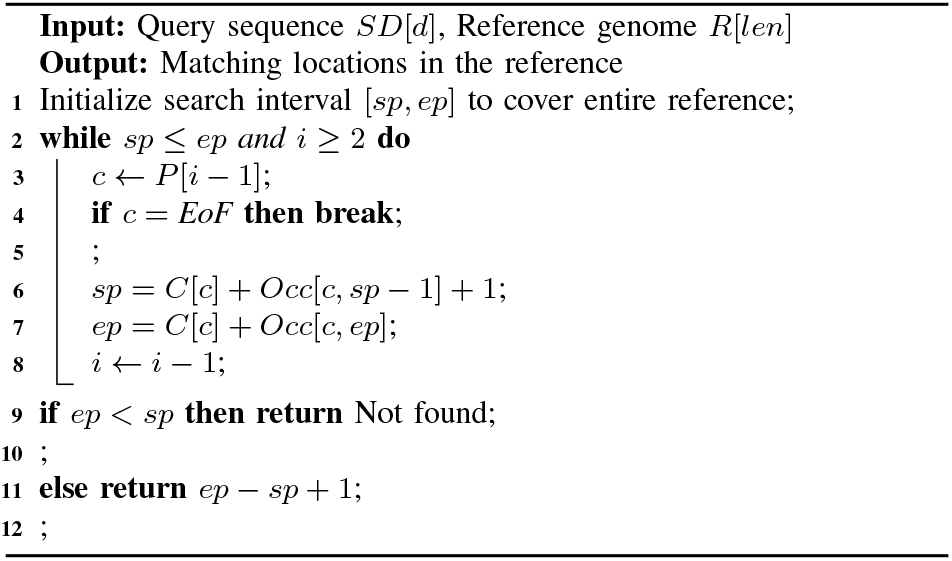
FM-index Based Seeding Algorithm

### B. Challenges of FM-index Search on Traditional Architectures

Despite the algorithmic elegance of FM-index, executing its search phase on conventional CPU-based systems presents significant performance challenges. Profiling studies (as illustrated in Figure 1) reveal that memory accesses dominate the overall execution cost. Specifically, dynamic random-access memory (DRAM) operations contribute approximately 60% to the cycles per instruction (CPI) overhead. Furthermore, load and store instructions constitute around 43.4% of the total executed instructions. The energy profile is similarly skewed, with nearly half (49.4%) of the total system energy consumed by DRAM accesses.

**Fig. 1.**
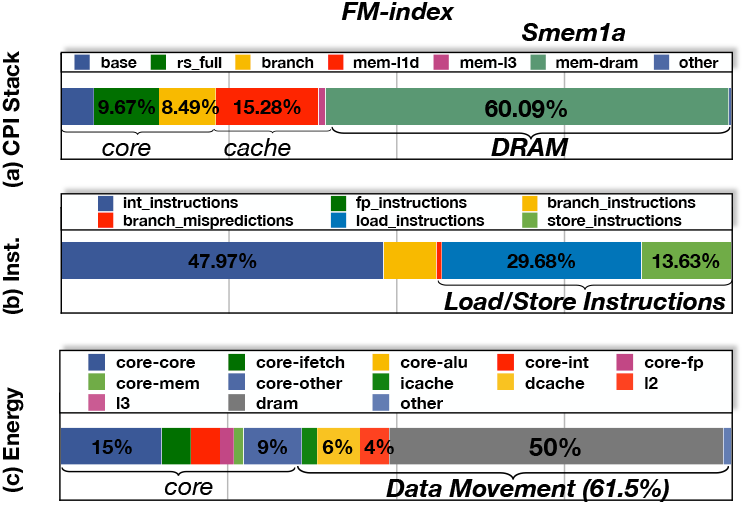
Profiling the FM-index kernel in BWA-MEM with Sniper and configuration in Table II: (a) CPI stack; (b) Instruction statistics; (c) Energy breakdown.

These observations highlight that the FM-index search is predominantly bottlenecked by memory bandwidth and latency, rather than computational complexity. The core reason lies in the nature of accessing the *Occ* data structure, which stores occurrence counts at multiple positions. For each base read, new search interval boundaries (*sp, ep*) must be calculated by retrieving values from *Occ* at locations dependent on the current interval. These accesses tend to be irregular and lack predictable spatial or temporal locality, resulting in extensive random memory accesses akin to pointer chasing.

Moreover, the FM-index search is inherently sequential since the interval computed at step *i* depends on the results from step *i*™1. This sequential dependency limits the opportunity for instruction-level parallelism (ILP) and diminishes the effectiveness of typical processor optimizations such as caching, prefetching, and multi-level memory hierarchies.

Previous research [8] has demonstrated that the computational demands of FM-index search are relatively simple, dominated by basic operations such as bitwise manipulation, integer addition and subtraction, and memory copy instructions. Complex operations like floating-point arithmetic, multiplication, division, or vectorized instructions are notably absent. Consequently, the performance bottleneck is primarily caused by the vast data movement rather than the arithmetic intensity. This leads to a pivotal question: *how can one supply the enormous memory bandwidth required by such lightweight computations efficiently?*

### C. Processing-In-Memory (PIM) as a Solution

Processing-in-memory (PIM) technology has emerged as a promising approach to address the bottlenecks posed by data movement in memory-intensive applications like FM-index search. The advent of 3D-stacked DRAM architectures has facilitated PIM by vertically integrating multiple DRAM layers with a logic layer using Through-Silicon Vias (TSVs), enabling unprecedented bandwidth between the memory and processing units.

Examples of such architectures include High Bandwidth Memory (HBM) [9] and Hybrid Memory Cube (HMC) [9], which embed a dedicated logic layer beneath multiple DRAM dies. Unlike traditional CPU systems constrained by limited off-chip memory bandwidth (e.g., 320 GB/s in a system with 16 8GB HMC modules), PIM can provide aggregate internal bandwidths as high as 8 TB/s by distributing computation close to or within the memory stack itself. By integrating compute units directly into or near the memory layers, PIM architectures can drastically reduce the costly data transfers between the CPU and main memory, significantly lowering latency and energy consumption. This is especially advantageous for FM-index search where the data access pattern is irregular and exhibits poor locality, rendering traditional caching and prefetching techniques ineffective.

Additionally, the independence of individual seed search tasks in FM-index makes the problem highly amenable to massive parallelism within PIM-enabled systems. However, designing effective PIM architectures for FM-index search involves overcoming two major challenges:

1. Architecting compute logic that can fully exploit the high internal bandwidth of 3D-stacked memory while remaining area- and energy-efficient.
2. Developing scalable communication protocols and data-sharing mechanisms between parallel processing elements embedded within the memory to coordinate and combine partial results efficiently.

This paper aims to explore these challenges and propose a novel PIM-based architecture tailored for FM-index search, enabling a breakthrough in performance and energy efficiency for genome sequence alignment workloads.

## III. GeneF Architecture

### A. Overview

Figure 2 depicts the conceptual architecture of GeneF. The system integrates 16 Hybrid Memory Cube (HMC) modules, collectively providing 128GB of memory capacity as shown in Figure 2(a). Each HMC cube, detailed in Figure 2(b), consists of eight stacked 8Gb DRAM layers and a logic layer. The cube is vertically partitioned into 32 Vaults, interconnected by an on-chip network.

**Fig. 2.**
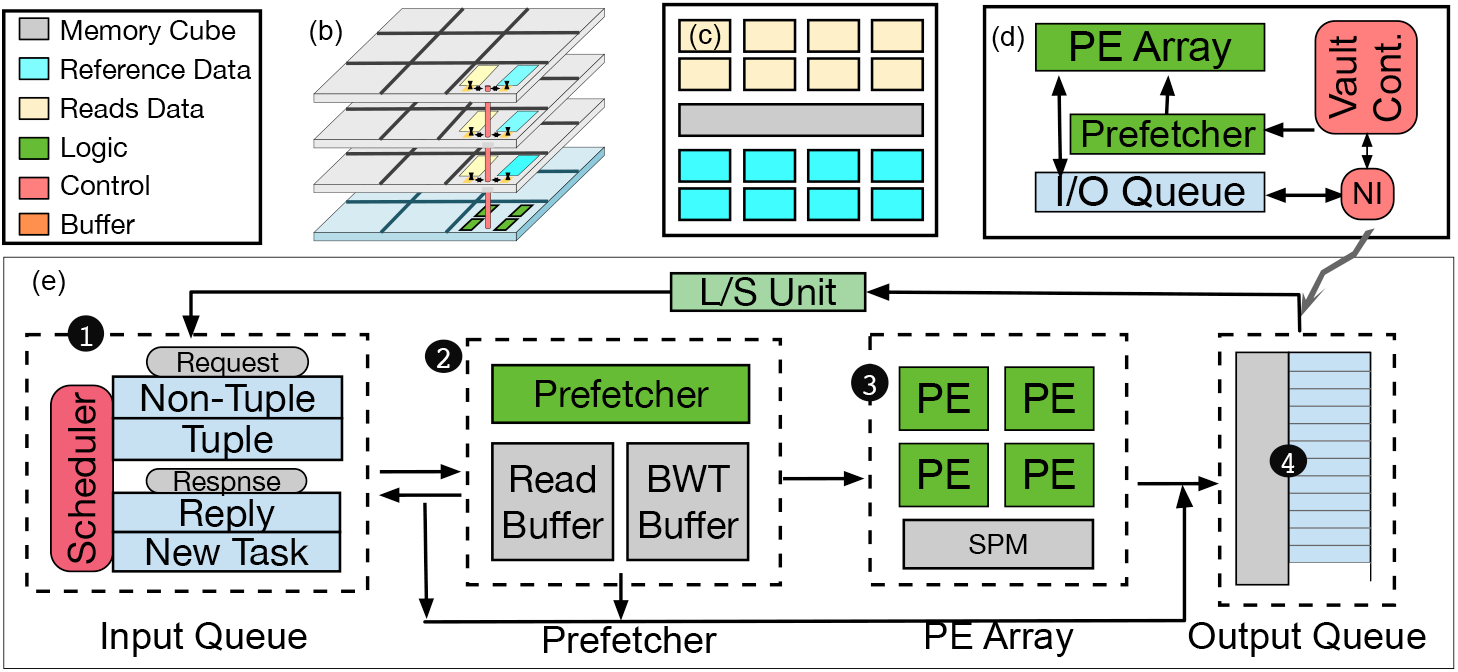
Architecure overview of GeneF accelerator (a) Network of Cubes, (b) Overview of a cube, (c) Data Mapping (d) Vault overview, (e) Compute Logic

These 32 Vaults per HMC are split into two groups (Figure 2(c)): one group stores reference genome data (blue), while the other manages DNA sequences (yellow). The logic layer structure, presented in Figure 2(d) and expanded in Figure 2(e), hosts custom RISC-V based processing elements (PEs), which are modular and replaceable. To facilitate communication between PEs across Vaults, a lightweight message passing protocol is implemented (Section III-B). The PEs focus solely on computation, with data-access duties delegated to a dedicated prefetcher (Section III-C). The Input Queue ❶ and Output Queue ❹ interface with the on-chip network, handling incoming requests and outgoing processing commands, respectively.

By independently processing reads and separating reference genome and DNA sequences into distinct Vault groups, the architecture enables parallel, independent execution within HMC cubes. This design achieves an internal bandwidth up to 8TB/s.

### B. Lightweight Message Passing Mechanism

Unlike host processors with full HMC address access, each GeneF core accesses only its local DRAM partition. Because reference genome queries involve random access across Vaults, remote data must be retrieved to process local requests.

Figure 4(a) illustrates the naive approach where a local PE sends a request to a remote Vault, waits for the data return, then resumes processing. This leads to idle time and high network traffic—e.g., FM-index’s *Occ4* requires 128 bytes per iteration, saturating the 16B/cycle network bandwidth [10].

To reduce overhead, computation migrates to the Vault containing the data. The lightweight message structure (Figure 3) comprises:

**Fig. 3.**
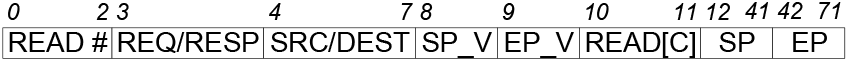
Lightweight Message Passing

- **READ#:** 3 bits to identify read slot (4–8 slots per Input Queue).
- **RES/RESP:** 1 bit indicating request or response.
- **SRC/DEST:** 4 bits each, source and destination Vault IDs (0– 15).
- **SP V/EP V:** Flags marking SP and EP validity.
- **READ[C]:** 2 bits representing current base.
- **SP/EP:** 30-bit indices for LF mapping.

Local Vaults send messages with LF mapping parameters to remote Vaults, which perform computation locally and return updated indices. This reduces network load to 72 bits per message versus 128 bytes, significantly improving throughput.

Figure 4(b) introduces a non-blocking remote function call: the source Vault sends a request then immediately frees its resources. Incoming requests are queued and scheduled for execution without stalling the source. This pipeline increases resource utilization and throughput.

**Fig. 4.**
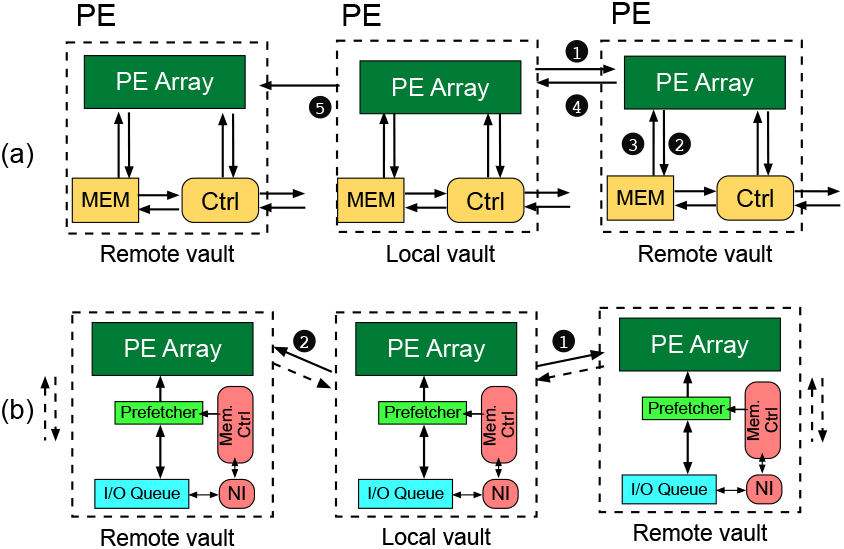
Non-Blocking Remote Function Call

#### a) Dynamic Scheduling Opportunities

To further optimize resource utilization, dynamic workload distribution strategies could be employed across Vaults. For example, adaptive task migration or workload-aware scheduling could balance memory pressure and prevent hotspots, especially in cases involving irregular or sparse query distributions.

### C. Prefetching

Prior PIM accelerators [8] have each PE complete LF mapping by issuing memory requests, integrating address translation (AU), memory access (MU), and calculation units (CU). This causes PE stalls during data fetch.

Although PIM’s proximity to memory yields high bandwidth, stalls occur due to remote Vault requests and limited logic area restricting PE count.

To mitigate this, we simplify PEs to only core computation and small registers, offloading address translation and memory access to a dedicated prefetcher. The scheduler translates addresses, dispatches them to the prefetcher, which loads data into a cache accessible by the PE array. This design pipelines fetch and compute stages, reducing stalls and maximizing throughput.

### D. Exploration of PE Count per Vault

With decoupled computation and memory access, a single PE per Vault underutilizes bandwidth. Multiple PEs improve utilization but must be balanced against data supply.

Figure 5(a) shows PE utilization versus PE count. Under three PEs, the aggregate compute rate cannot consume prefetch data, causing underutilization despite 100% PE utilization individually. At four PEs, the system saturates bandwidth, and idle rates drop to 10–20%, indicating optimal balance.

**Fig. 5.**
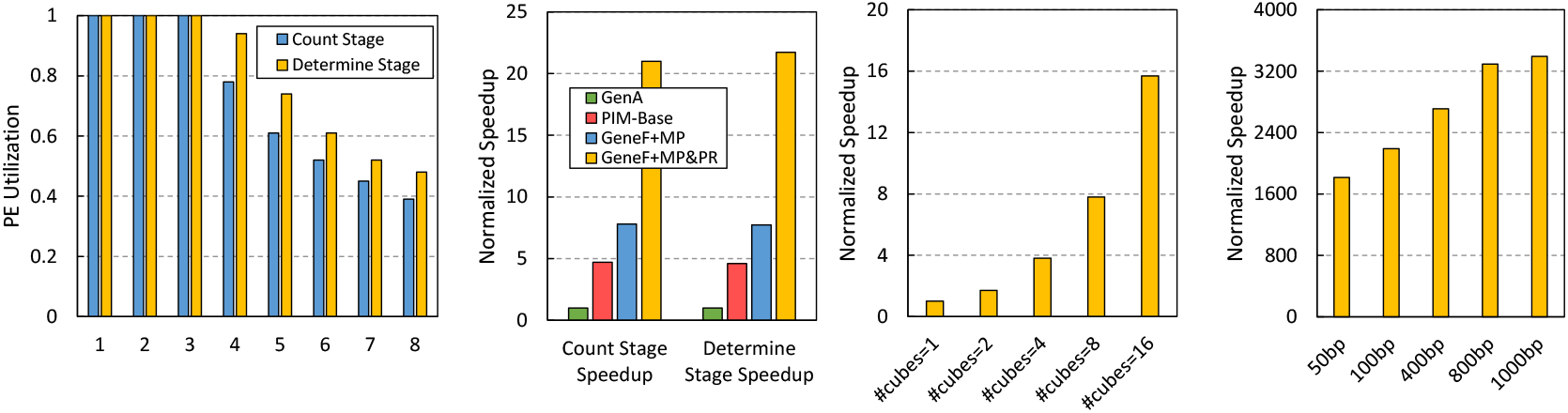
Profiling the FM-index kernel in BWA-MEM with Sniper and configuration in Table II: (a) CPI stack; (b) Instruction statistics; (c) Energy breakdown.

More than four PEs cause higher idle times as computation outpaces data supply, reducing efficiency. We select four PEs per Vault as the optimal configuration balancing throughput and resource use.

## IV. Experimental Evaluation

### A. Experimental Setup

To evaluate the effectiveness of our proposed GeneF architecture, we conducted comprehensive experiments focusing on its performance, scalability, and energy efficiency. We compared GeneF against several baseline systems including a 32-thread CPU implementation and the Niubility accelerator—a specialized FM-index computing logic—which serves as a representative state-of-the-art prior work. Our experiments simulate real-world sequencing workloads, particularly targeting typical next-generation sequencing (NGS) read lengths. Performance metrics such as throughput, speedup, and energy consumption were collected and analyzed to demonstrate the benefits of our design choices.

### B. Performance Analysis

Figure 5(b) presents a detailed comparison of the performance achieved by GeneF against the baseline architectures. We normalized the performance metric with respect to Niubility, setting its throughput to 1. Niubility employs a custom FM-index logic and achieves a 79× speedup compared to a 32-thread CPU baseline. Building upon Niubility’s design, GeneF integrates a 3D-stacked memory architecture to exploit higher memory bandwidth and parallelism.

Our results show that GeneF attains approximately a 20× speedup over a corresponding ASIC implementation. When compared to the CPU baseline, the *count* and *determine* stages of the FM-index algorithm accelerate by factors of 1820× and 1728×, respectively. This significant acceleration stems from both the superior memory band- width enabled by the Hybrid Memory Cube (HMC) and optimizations tailored to the novel memory structure. Key among these is the partitioning of reference genome data and grouping within memory regions to optimize data locality. Furthermore, the implementation of a lightweight, non-blocking message-passing mechanism combined with remote in-place computation effectively hides the latency inherent in cross-vault communication. The decoupling of computation from memory access, along with proactive data prefetching, further enhances the efficiency and resource utilization, culminating in the observed speedups.

#### a) Extended Comparisons

While Niubility and Darwin represent strong baselines, other accelerators like GenAx and AIM offer alternative design philosophies. While direct comparisons are challenging due to architectural and algorithmic differences, future work could implement or simulate these baselines within a unified framework to quantify trade-offs in throughput, energy, and scalability more comprehensively.

### C. Efficiency of Message Passing and Prefetch Mechanisms

While near-memory computing inherently offers higher bandwidth to support massive parallelism, simply scaling existing accelerators to 3D-stacked memory architectures does not guarantee ideal performance. As shown in Figure 5(b), scaling the number of processing elements (PEs) from 384 to 2048 yields only a 4.7× and 4.6× speedup for the *count* and *determine* stages, respectively. This sublinear scaling occurs because, although memory bandwidth increases, the HMC’s capacity constraints require dispersing reference data across multiple vaults, introducing latency from remote vault data access.

Our solution to this challenge is a carefully designed lightweight message-passing protocol tailored to the application’s characteristics (discussed in Section III). This mechanism effectively masks the overhead caused by remote vault communication and balances the workload pressure. Compared to a baseline PIM design without message passing (PIM-Base), our message mechanism improves performance by 1.66× and 1.68× in the *count* and *determine* stages, respectively.

In addition, to address the logic layer area and power consumption limitations inherent in 3D-stacked memory systems, we minimized the PE design and separated computation from memory access. Utilizing a dedicated prefetcher, data is loaded into local caches before computation begins, ensuring PEs are never starved for data. This prefetching approach further boosts performance by 2.69× in the *count stage* and 2.81× in the *determine stage*.

### D. Scalability Study

To assess GeneF’s scalability, we increased memory capacity by adding more HMC cubes and examined how performance scales with parallelism and workload distribution. While increasing the number of PEs generally improves parallelism, it can also reduce workload locality per PE, potentially degrading performance due to increased data access overheads.

Figure 5(c) illustrates GeneF’s speedup as a function of the number of HMC cubes. Using two cubes results in a 1.9× performance gain, while scaling up to sixteen cubes achieves an average speedup of 15.7× compared to the baseline single-cube setup. This near-linear scalability arises from the intrinsic properties of the BWT-based gene alignment algorithm, which exhibits natural concurrency due to the independence of individual read alignments. The relatively small memory footprint of the BWT structure allows the reference and read data to be partitioned without requiring inter-cube communication, enabling independent processing across multiple HMC units. Consequently, the HMC architecture effectively supports horizontal scaling with minimal communication overhead, demonstrating GeneF’s extensibility.

### E. Impact of Read Length on Performance

GeneF targets acceleration of FM-index algorithms widely used in genomic applications like BWA-MEM, which combines FM-index-based seed matching with dynamic programming-based seed extension (e.g., Smith-Waterman). To understand GeneF’s performance implications in practical pipelines, we analyzed its impact on BWA-MEM using Darwin [11], a state-of-the-art hardware accelerator for the Smith-Waterman algorithm, as a comparison baseline.

As illustrated in Figure 5(d), accelerating only the dynamic programming component yields a modest 2.2× speedup on a 16-thread CPU due to the significant proportion of execution time consumed by the FM-index phase. Incorporating GeneF to accelerate the FM-index stage, alongside Darwin’s Smith-Waterman accelerator, results in a dramatic improvement exceeding 1000× speedup for reads of length 101 base pairs, typical of NGS technologies [12], [13].

GeneF is compatible with varying read lengths and continues to offer increasing speedups for longer reads, as shown in Figure 5(d). Longer reads increase memory traffic, which aligns well with the strengths of near-data processing. Since ultra-long reads remain a memory-bound challenge in sequence alignment, we anticipate that GeneF will maintain or even improve performance benefits as read lengths increase further. To further validate GeneF’s generalizability, ongoing work involves evaluating its performance on a broader spectrum of real-world genomic datasets with variable characteristics, such as different species, mutation rates, and sequencing depths. This would demonstrate the robustness of the architecture across diverse sequencing applications beyond typical NGS pipelines.

### F. Energy and Power Efficiency

Energy consumption is a critical consideration for near-memory accelerators. Figure 6 presents the normalized energy usage of various HMC-based systems, including GeneF. Our power model, detailed in Section 5, accounts for both logic and memory layers, as well as PE activity.

**Fig. 6.**
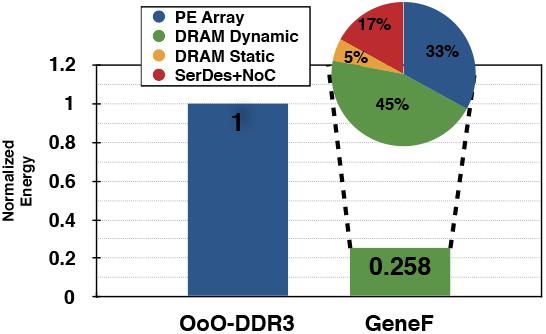
GeneP area and power consumption

GeneF consumes only about 25% of the energy required by conventional CPU-DDR3 systems using out-of-order cores. Within GeneF, power consumption is almost evenly split between logic and memory layers. The dynamic power of DRAM, including data accesses and row activations, dominates the memory power profile, while static power accounts for only 5% of the total energy, indicating highly efficient bandwidth utilization.

The SerDes circuits responsible for off-chip communication contribute approximately 17% of GeneF’s power consumption, whereas the PEs account for 33%. Among PE components, the on-chip network is the largest consumer of power, while SerDes power remains low due to minimal off-chip communication. This balance demonstrates effective utilization of computational resources and communication channels.

The logic layer energy density of GeneF stands at 20.7 mW/mm^2^, well below the thermal design limit of 133 mW/mm^2^. The total PE logic occupies roughly 5.1 mm^2^, which is only 2.3% of the logic layer’s area, indicating substantial headroom for future expansion. Overall, GeneF is thermally sustainable and significantly reduces energy consumption, achieving energy efficiency improvements of 7032× and 6687× in the *count* and *determine* stages, respectively.

### Discussion on Limitations

While GeneF demonstrates excellent performance and energy efficiency, practical deployment may face hardware-level constraints. Notably, the logic layer’s thermal density, though below current design limits, could pose scalability concerns under sustained highthroughput workloads. Further, under scenarios where memory pressure is low, processing elements (PEs) may face underutilization, leading to suboptimal resource efficiency. Future designs could explore adaptive scaling techniques and thermal-aware scheduling to mitigate such risks.

### Hardware Cost and Scalability Considerations

From a deployment standpoint, the scalability of GeneF must consider manufacturing and economic factors. The use of 3D-stacked memory, while powerful, introduces fabrication complexity and cost. However, the demonstrated area efficiency (only 2.3% of logic layer) and modest power density suggest that scaling GeneF to larger systems is feasible without prohibitive cost increases. Incorporating commodity 3D memory with programmable logic could offer costeffective entry points for practical adoption.

## V. Conclusion

In this work, we conduct a thorough investigation of the computational and memory access patterns inherent to the FM-index algorithm, which is foundational to many genomic data processing tasks. Our detailed analysis highlights that a substantial portion of the total system energy consumption arises from the frequent data movement between memory and compute units, rather than from the computational operations themselves. Specifically, this energy overhead is largely driven by several simple yet repetitive data manipulation functions.

Recognizing these challenges, we revisit the concept of processingin-memory (PIM) in the modern context, motivated by two key technological trends. First, advancements in 3D-stacked memory technology enable the close integration of logic and memory layers, reducing latency and increasing bandwidth significantly. Second, contemporary genomic processing workloads demand massive memory bandwidth and are highly data-intensive, making them ideal candidates for PIM-based acceleration.

To address these requirements, we propose *GeneF*, a novel PIM accelerator tailored for DNA sequence alignment tasks. *GeneF* integrates multiple lightweight, RISC-V inspired processing elements (PEs) directly within the logic layer of 3D-stacked memory. To overcome latency issues associated with remote memory accesses, we design an efficient, lightweight messaging mechanism that leverages in-place computation to mask communication delays. Furthermore, to simplify PE design and maximize utilization, we introduce specialized prefetching strategies that proactively prepare data for computation.

Our evaluation demonstrates that *GeneF* delivers substantial improvements over conventional CPU platforms and even custom ASIC solutions, achieving significant gains in both computational speed and energy efficiency. These results underscore the potential of reducing costly data transfers through tightly coupled computation and memory as a promising direction for future high-performance genomic data processing. We believe that the principles embodied in *GeneF* can inspire the design of next-generation accelerators that meet the growing demands of genomics and other data-centric workloads.

## References

[1] J. Shendure and H. Ji, “Next-generation DNA sequencing,” Nature biotechnology, vol. 26, no. 10, pp. 1135–1145, 2008.

[2] M.-C. F. Chang, Y.-T. Chen, J. Cong, P.-T. Huang, C.-L. Kuo, and C. H. Yu, “The smem seeding acceleration for dna sequence alignment,” in Field-Programmable Custom Computing Machines (FCCM), 2016 IEEE 24th Annual International Symposium on, pp. 32–39, IEEE, 2016.

[3] J. Cong, Z. Fang, M. Gill, F. Javadi, and G. Reinman, “AIM: accelerating computational genomics through scalable and noninvasive accelerator-interposed memory,” in Proceedings of the International Symposium on Memory Systems, pp. 3–14, ACM, 2017.

[4] Z. D. Stephens, S. Y. Lee, F. Faghri, R. H. Campbell, C. Zhai, M. J. Efron, R. Iyer, M. C. Schatz, S. Sinha, and G. E. Robinson, “Big data: astronomical or genomical?,” PLoS biology, vol. 13, no. 7, p. e1002195, 2015.

[5] H. Li, “Aligning sequence reads, clone sequences and assembly contigs with bwa-mem,” vol. 1303, 2013.

[6] B. Langmead, C. Trapnell, M. Pop, S. L. Salzberg, et al., “Ultrafast and memory-efficient alignment of short dna sequences to the human genome,” Genome biol, vol. 10, no. 3, p. R25, 2009.

[7] N. Ahmed, K. Bertels, and Z. Al-Ars, “A comparison of seed-and-extend techniques in modern DNA read alignment algorithms,” in Bioinformatics and Biomedicine (BIBM), 2016 IEEE International Conference on, pp. 1421–1428, IEEE, 2016.

[8] Y. Wang, X. Li, D. Zang, G. Tan, and N. Sun, “Accelerating fm-index search for genomic data processing,” in Proceedings of the 47th International Conference on Parallel Processing, ICPP 2018, (New York, NY, USA), pp. 65:1–65:12, ACM, 2018.

[9] C. Weis, N. Wehn, L. Igor, and L. Benini, “Design space exploration for 3d-stacked drams,” in 2011 Design, Automation & Test in Europe, pp. 1–6, IEEE, 2011.

[10] M. Gao, G. Ayers, and C. Kozyrakis, “Practical near-data processing for in-memory analytics frameworks,” in 2015 International Conference on Parallel Architecture and Compilation (PACT), pp. 113–124, IEEE, 2015.

[11] Y. Turakhia, G. Bejerano, and W. J. Dally, “Darwin: A genomics coprocessor provides up to 15,000 x acceleration on long read assembly,” in Proceedings of the Twenty-Third International Conference on Architectural Support for Programming Languages and Operating Systems, pp. 199–213, ACM, 2018.

[12] M. A. DePristo, E. Banks, R. Poplin, K. V. Garimella, J. R. Maguire, C. Hartl, A. A. Philippakis, G. Del Angel, M. A. Rivas, M. Hanna, et al., “A framework for variation discovery and genotyping using next-generation DNA sequencing data,” Nature genetics, vol. 43, no. 5, p. 491, 2011.

[13] D. Fujiki, A. Subramaniyan, T. Zhang, Y. Zeng, R. Das, D. Blaauw, and S. Narayanasamy, “GenAx: a genome sequencing accelerator,” in Proceedings of the 45th Annual International Symposium on Computer Architecture, pp. 69–82, IEEE Press, 2018.

